# The pathogenic roles of the p.R130S prestin variant in DFNB61 hearing loss

**DOI:** 10.1101/2023.08.21.554157

**Authors:** Satoe Takahashi, Yingjie Zhou, Mary Ann Cheatham, Kazuaki Homma

**Author notes:** Address correspondence to: Kazuaki Homma, 303 E Chicago Ave, Chicago, IL 60611. **Author Contributions:** K.H. initiated the study. S.T., Y.Z., M.A.C., and K.H. performed experiments and analyzed the data. S.T. and K.H. wrote the manuscript with input from the other authors.

## Abstract

DFNB61 is a recessively inherited nonsyndromic hearing loss caused by mutations in *SLC26A5*, the gene that encodes the voltage-driven motor protein, prestin. Prestin is abundantly expressed in the auditory outer hair cells that mediate cochlear amplification. Two DFNB61-associated *SLC26A5* variants, p.W70X and p.R130S, were identified in patients who are compound heterozygous for these nonsense and missense changes (*SLC26A5*^*W70X/R130S*^). Our recent study showed that mice homozygous for p.R130S (*Slc26a5*^*R130S/R130S*^) suffer from hearing loss that is ascribed to significantly reduced motor kinetics of prestin. Given that W70X-prestin is nonfunctional, compound heterozygous *Slc26a5*^*R130S/-*^ mice were used as a model for human *SLC26A5*^*W70X/R130S*^. By examining the pathophysiological consequences of p.R130S prestin when it is the sole allele for prestin protein production, we determined that this missense change results in progressive outer hair cell loss in addition to its effects on prestin’s motor action. Thus, this study fully defines the pathogenic roles for the p.R130S prestin, which points to the presence of a limited time window for potential clinical intervention.

## INTRODUCTION

*SLC26A5* encodes a membrane-based voltage-driven motor protein, prestin, that is comprised of 744 amino acids. Prestin is abundantly expressed in the lateral membrane of the auditory outer hair cells (OHCs) and confers voltage-dependent somatic elongation and contraction, referred to as electromotility (Brownell et al., 1985). This prestin-mediated OHC electromotility is thought to work against the viscous damping of the cochlear fluid to augment sound-induced motions of the organ of Corti, so that the inner hair cells (IHCs), the true sensory receptor cells in the cochlea, can respond to very low sound pressures, i.e., as low as ∼20 μPa. The indispensability of prestin for this mechanical augmentation process, i.e., cochlear amplification, has been experimentally demonstrated using mouse models lacking prestin (Cheatham et al., 2004; Cheatham et al., 2007; Liberman et al., 2002; Wu et al., 2004) and expressing a dysfunctional prestin mutant (Dallos et al., 2008).

Several *SLC26A5* variants are known to associate with nonsyndromic hearing loss, DFNB61 (OMIM: 613865). IVS2-2A>G is the earliest reported *SLC26A5* variant, which was presumed to compromise prestin protein production by disrupting pre-mRNA splicing (Liu et al., 2003). However, later studies questioned the pathogenicity of this variant (Shearer et al., 2014; Tang et al., 2005; Teek et al., 2009). Although it was experimentally demonstrated that this intronic mutation affects splicing of prestin mRNA in mice, prestin protein production remained unaffected, resulting in normal hearing (Zhang et al., 2016). Subsequently, a heterozygous missense variant, c.449G>A (p.R150Q), was identified in a hearing-impaired patient and his normal-hearing father (Toth et al., 2007). This missense change shifts the voltage operating point (V_pk_) of prestin to the hyperpolarizing direction, but the V_pk_ change is small when coexpressed with wildtype (WT) prestin (Toth et al., 2007). In any case, p.R150Q is unlikely pathogenic even in homozygotes, because mice expressing a manmade prestin mutant, “C1” (Oliver et al., 2001), whose V_pk_ is more negatively shifted compared to that of p.R150Q prestin, retain WT-like hearing (Gao et al., 2007). The p.W70X (c.209G>A) and p.R130S (c.390A>C) *SLC26A5* variants were identified in two young patients with moderate to profound hearing loss (Matsunaga and Morimoto, 2016; Mutai et al., 2013). Both patients were compound heterozygous for these two prestin variants, whereas their normal-hearing parents were heterozygous for each variant (*SLC26A5*^*W70X/+*^ and *SLC26A5*^*R130S/+*^), which is in line with the recessive nature of DFNB61. Since p.W70X terminates translation of prestin protein before the first transmembrane domain, it is likely that this nonsense change results in a prestin-knockout (KO)-like null *SLC26A5* allele. The p.R130S variant significantly slows prestin’s motor kinetics (Takahashi et al., 2016a) and results in hearing loss in R130S-prestin mouse models (Takahashi et al., 2023). Collectively, it is likely that compound heterozygosity for p.W70X and p.R130S *SLC26A5* variants is causally associated with DNB61 hearing loss. Given the recessive inheritance of the pathogenic prestin variants, one promising clinical solution for DFNB61 hearing loss is to introduce WT prestin-encoding cDNA to affected OHCs. However, such a genetic or any other clinical intervention is based on the premise that the target cell type, i.e., OHC, is remained in the cochlea.

OHCs lacking prestin (prestin-KO) are shorter and exhibit a reduced axial stiffness compared to WT. They also suffer from progressive degeneration (Cheatham et al., 2007; Dallos et al., 2008; Liberman et al., 2002; Wu et al., 2004), implying that prestin is essential to the OHC’s structural and mechanical integrity. Since p.R130S partially impairs membrane targeting of the prestin protein (Takahashi et al., 2016a; Takahashi et al., 2023), it is conceivable that OHC integrity is impaired in patients who carry only one copy of p.R130S-prestin. The objective of this study was to define the pathophysiological consequences of having only one p.R130S prestin allele. We found that W70X-prestin is nonfunctional and thus used *Slc26a5*^*R130S/-*^ mice as a model for human *SLC26A5*^*W70X/R130S*^. This compound heterozygous mouse model expresses significantly reduced R130S-prestin protein compared to homozygotes (*Slc26a5*^*R130S/R130S*^) and exhibits a more severe, progressive OHC loss. This study thus defines an additional pathogenic role for the p.R130S prestin variant and indicates the presence of a limited therapeutic time window for potential clinical interventions targeting the affected OHCs.

## RESULTS

### OHC motor function is significantly attenuated in *Slc26a5*^*R130S/-*^ compound heterozygotes

In OHCs the nonsense-mediated mRNA decay mechanism likely destroys prestin mRNA with c.209G>A (p.W70X) because this point mutation results in a premature stop codon >80 nucleotides upstream of the 3’ end of exon 4. Even if this cellular surveillance mechanism did not work perfectly, the nonsense change terminates translation of prestin protein before the first transmembrane domain (**Fig. 1A**) and thus would not allow production of a functional motor protein. In HEK293T cells, we experimentally confirmed that W70X-prestin is retained intracellularly (**Figs. 1B and 1C**) and does not confer nonlinear capacitance (NLC), which is the electrical signature of electromotility used as a proxy for evaluating prestin’s motor activity (Ashmore, 1990; Santos-Sacchi, 1991) (**Fig. 1D**). Thus, the p.W70X prestin mutation likely results in a null allele.

**Fig. 1.**
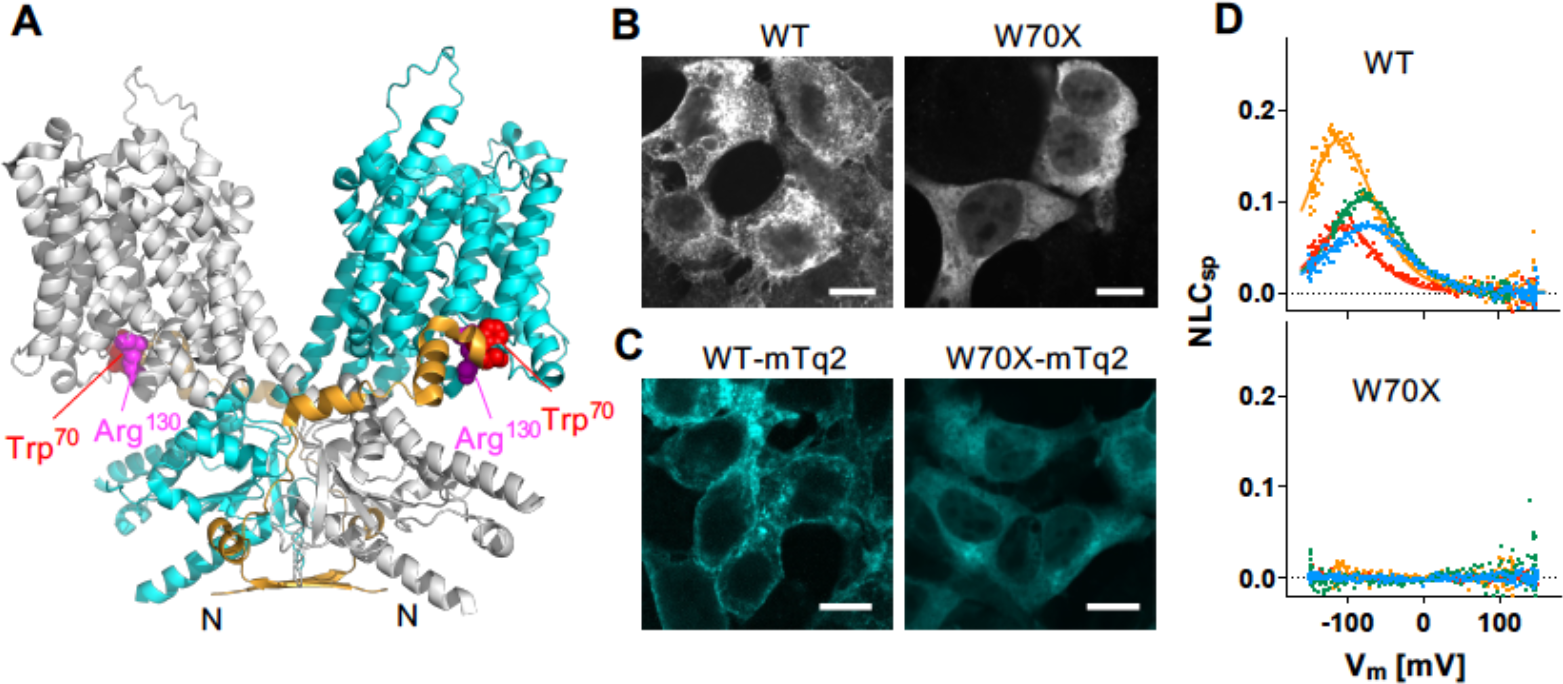
W70X-prestin is nonfunctional. (**A**) Prestin homodimer (PDB: 7LGU) showing one protomer in gray and the other in cyan, with the N-terminal amino acid residues Thr^11^-Lys^69^ shown in orange. The Trp^70^ and Arg^130^ residues are shown with a sphere representation in red and magenta, respectively. N, N-terminus. (**B**) Representative fluorescence images of HEK293T cells expressing WT-or W70X-prestin, stained with anti-N-terminal prestin antisera. Scale bars, 10 μm. (**C**) Representative fluorescence images of HEK293T cells expressing WT-or W70X-prestin with a C-terminally attached mTurquoise2 (mTq2) tag. Scale bars, 10 μm. (**D**) NLC recorded in HEK293T cells expressing WT-or W70X-prestin. Four examples are shown in different colors for each. In WT, the solid lines indicate two-state Boltzmann curve fits. The magnitude of NLC (C_m_ - C_lin_) was corrected for the cell size (C_lin_), i.e., NLC_sp_ ≡ (C_m_ - C_lin_)/C_lin_.

To understand the pathology of having only one copy of the p.R130S *SLC26A5* allele as in human patients, we generated *Slc26a5*^*R130S/-*^ compound heterozygous mice by crossing R130S-prestin mice (Takahashi et al., 2023) with prestin-KO mice (Cheatham et al., 2005; Liberman et al., 2002). Our preceding studies (Takahashi et al., 2016a; Takahashi et al., 2023) showed that the p.R130S missense change slows the voltage-driven motor kinetics by reducing the binding affinity of prestin to its extrinsic cofactor, chloride, which is essential for prestin’s fast motor action. These studies also showed that the p.R130S missense change partially impairs membrane targeting. In OHCs of *Slc26a5*^*R130S/R130S*^ homozygotes, the partially impaired membrane targeting of the R130S-prestin protein was quantitatively reflected in reductions of the linear and nonlinear cell membrane electric capacitance (C_lin_ and NLC, respectively) (Takahashi et al., 2023). Thus, it is conceivable that expression of R130S-prestin is reduced to a greater degree in OHCs of *Slc26a5*^*R130S/-*^ compound heterozygotes, which might result in a more severe functional defect in OHCs. We explored this possibility by measuring the cell membrane electric capacitance, C_m_ =NLC + C_lin_, in OHCs isolated from *Slc26a5*^*R130S/-*^ compound heterozygous mice. Since previous work indicated that OHCs isolated from *Slc26a5*^*+/-*^ heterozygous mice have near-normal hearing (Cheatham et al., 2007; Cheatham et al., 2005; Liberman et al., 2002), they were also included as positive controls. Representative C_m_ data and summaries of the NLC and C_lin_ parameters are provided in **Figs. 2A and 2B**, respectively. As anticipated, significant reductions of NLC (NLC_pk_) and C_lin_ were detected in *Slc26a5*^*R130S/-*^ compared to WT controls (**Fig. 2B**). Based on our recent study (Takahashi et al., 2023), the NLC_pk_ and C_lin_ values of *Slc26a5*^*R130S/-*^ were smaller than in *Slc26a5*^*R130S/R130S*^ homozygotes. Comparisons between *Slc26a5*^*R130S/-*^ and *Slc26a5*^*R130S/R130S*^ are as follows: NLC_pk_: 1.1 ± 0.4 pF (mean ± SD, n=60) vs. 3.2 ± 0.7 pF (mean ± SD, n=94), *p* < 0.0001; C_lin_: 5.0 ± 0.6 pF (mean ± SD, n=60) vs. 5.4 ± 0.5 pF (mean ± SD, n=94), *p* < 0.0001), indicating reduced expression of R130S-prestin in the lateral membrane of OHCs in *Slc26a5*^*R130S/-*^ compound heterozygotes.

**Fig. 2.**
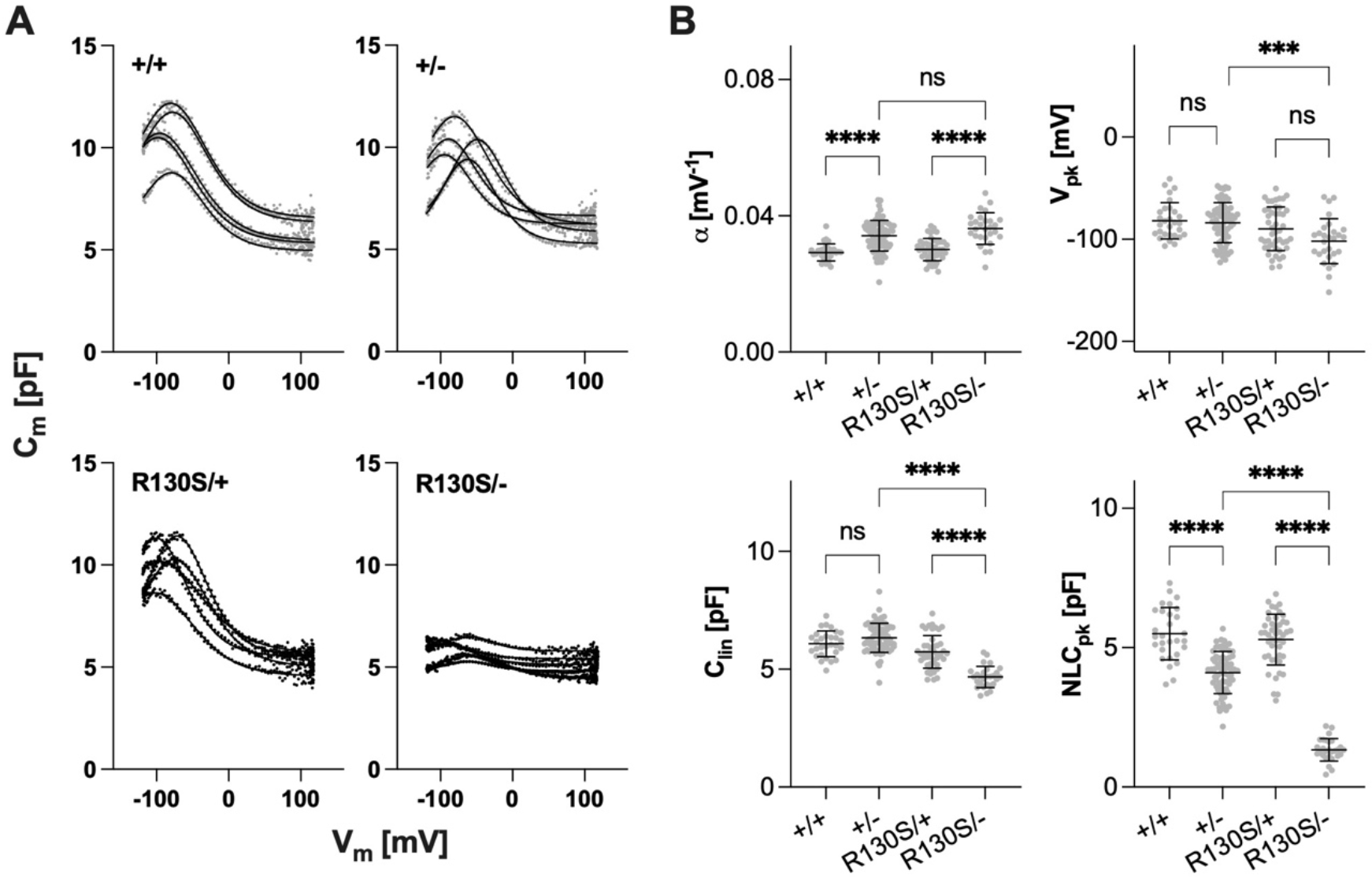
NLC recordings in OHCs. (**A**) Examples of NLC measurements. Solid lines indicate two-state Boltzmann curve fits. (**B**) Summaries of NLC parameters. α is the slope factor of the voltage-dependence of prestin-mediated charge transfer, V_pk_ is the voltage at which the maximum charge movement is attained, C_lin_ is the linear capacitance, and NLC_pk_ is the magnitude of NLC at V_pk_. Horizontal solid lines indicate the means and standard deviations. OHCs were collected from the apical region (corresponding to 4-10 kHz) of the cochlea (P25-33). The *p* values were computed by One-way ANOVA combined with the Tukey’s post hoc test. “ns”, *p* ≥ 0.05. “^***^”, 0.001 ≤ *p* < 0.0001. “^****^”, *p* < 0.0001.

### Distortion product otoacoustic emissions are vastly reduced in *Slc26a5*^*R130S/-*^ compound heterozygotes

To determine the physiological consequences of the reduced expression of R130S-prestin in *Slc26a5*^*R130S/-*^, we measured distortion product otoacoustic emissions (DPOAEs) that are associated with OHC mechanoelectrical transduction, a nonlinear process. Hence, DPOAEs are used to assess OHC function *in vivo*. In this procedure, two pure tones are simultaneously presented at f_1_ and f_2_ with a fixed frequency ratio, f_2_/f_1_ = 1.2, and at equal levels (L_1_= L_2_= 70 dB SPL). These iso-input functions or DP grams revealed that *Slc26a5*^*R130S/-*^ compound heterozygous mice had vastly reduced DPOAEs at 2f_1_-f_2_ compared to *Slc26a5*^*+/+*^ (black) and *Slc26a5*^*+/-*^ (blue) mice (**Fig. 3**). While responses were relatively higher at or a few days before weaning (P17-21, magenta), the DPOAEs at 70 dB SPL for mice at one month (P27-33, brown) and at six weeks of age (P42-43, green) were near or within the noise floor (gray).

**Fig. 3.**
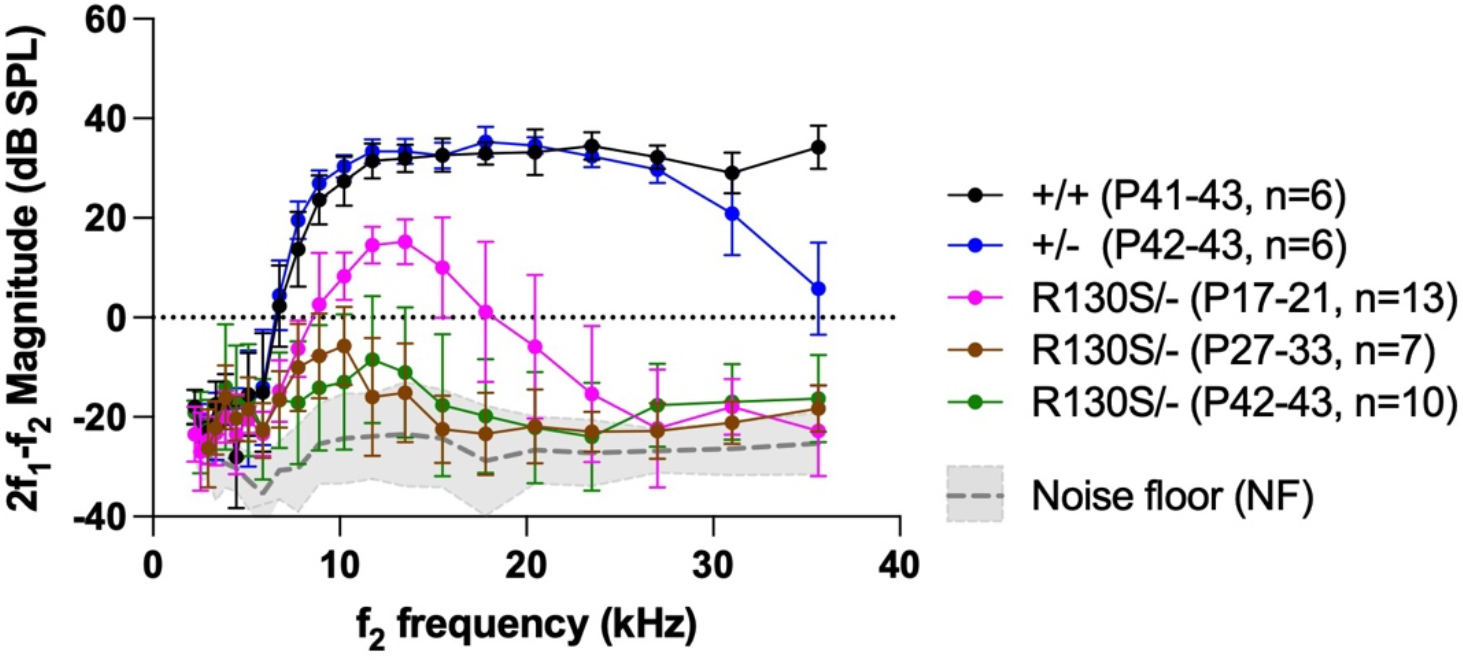
DPOAE iso-input functions. DPOAEs of *Slc26a5*^*R130S/-*^ (R130S/-) mice tested at P17-21 (magenta), P27-33 (brown), and P42-43 (green) were shown together with those of *Slc26a5*^*+/+*^ (black) and *Slc26a5*^*+/-*^ (blue) measured at P41-42. The average noise floor is plotted in gray dotted line with standard deviations colored in gray. The sound pressure levels (SPL, L_1_ and L_2_) at f_1_ and f_2_ frequencies (f_2_/f_1_=1.2) were both 70 dB SPL. Horizontal solid lines indicate the means and standard deviations.

To determine if DPOAEs were generated at higher levels, input-output or growth functions were also collected with f_2_ frequencies at 8, 12, 27, and 32 kHz. Although f_2_/f_1_ remained fixed at 1.2, the levels of f_1_ were 10 dB greater than those of f_2_ (L_1_ = L_2_+10 dB) (**Fig. 4A**). The functions in *Slc26a5*^*R130S/-*^ compound heterozygous mice are shifted horizontally to the right indicating elevation in DPOAE thresholds, the level of f_1_ generating a DPOAE of 0 dB SPL. It is notable that only the very youngest mice at P17-21 show growth of 2f_1_-f_2_ that is distinct from the coupler measurements plotted with gray dotted lines. For older mice and for all mutants at f_2_=27 and 32 kHz, the DPOAE magnitudes are small and similar to the values measured in the coupler. Hence, the DPOAE thresholds plotted in **Fig. 4B** relate to OHC function but only for the youngest *Slc26a5*^*R130S/-*^ mice and only at f_2_=8 and 12 kHz. All other “thresholds” can be explained by distortion in the sound as indicated by the upward pointing arrows. Because the stimulus itself is nonlinear at high stimulus levels, there is energy at 2f_1_-f_2_ that is picked up by the microphone in the ear canal. In **Fig. 4C**, the DPOAE threshold shifts are plotted for different ages. In this plot, the shifts are determined by comparing the thresholds in age-matched controls. In other words, the threshold shifts are computed as the difference between age-matched controls and mutants at P17-21, P27-33, and P41-43. Again, the upward pointing arrows indicate that the “threshold” cannot be determined because of distortion in the sound. The magnitude of NLC at V_pk_ (NLC_pk_) is also plotted in **Fig. 4D** as a function of age, which refutes the possibility that the larger DPOAEs in young *Slc26a5*^*R130S/-*^ mice may be ascribed to greater expression of prestin proteins. We presume that larger DPOAEs in *Slc26a5*^*R130S/-*^ mice at or before weaning relate to the relatively higher chloride concentrations in immature OHCs (Takahashi et al., 2023). Taken together, the progressive and frequency-dependent DPOAE threshold shifts found in *Slc26a5*^*R130S/-*^ mice reaffirms the pathogenicity of p.R130S prestin, consistent with its role in slowing prestin’s motor activity as demonstrated in our recent study (Takahashi et al., 2023). It is intriguing that DPOAE thresholds were similar for *Slc26a5*^*R130S/-*^ compound heterozygotes and *Slc26a5*^*R130S/R130S*^ homozygotes despite the fact that R130S-prestin expression is much lower in *Slc26a5*^*R130S/-*^ compared to *Slc26a5*^*R130S/R130S*^. This observation is evocative of the hypothetical turgor pressure-powered electromotility mechanism (Homma et al., 2021), which is more sensitive to prestin’s motor kinetics than to the amount of prestin protein.

**Fig. 4.**
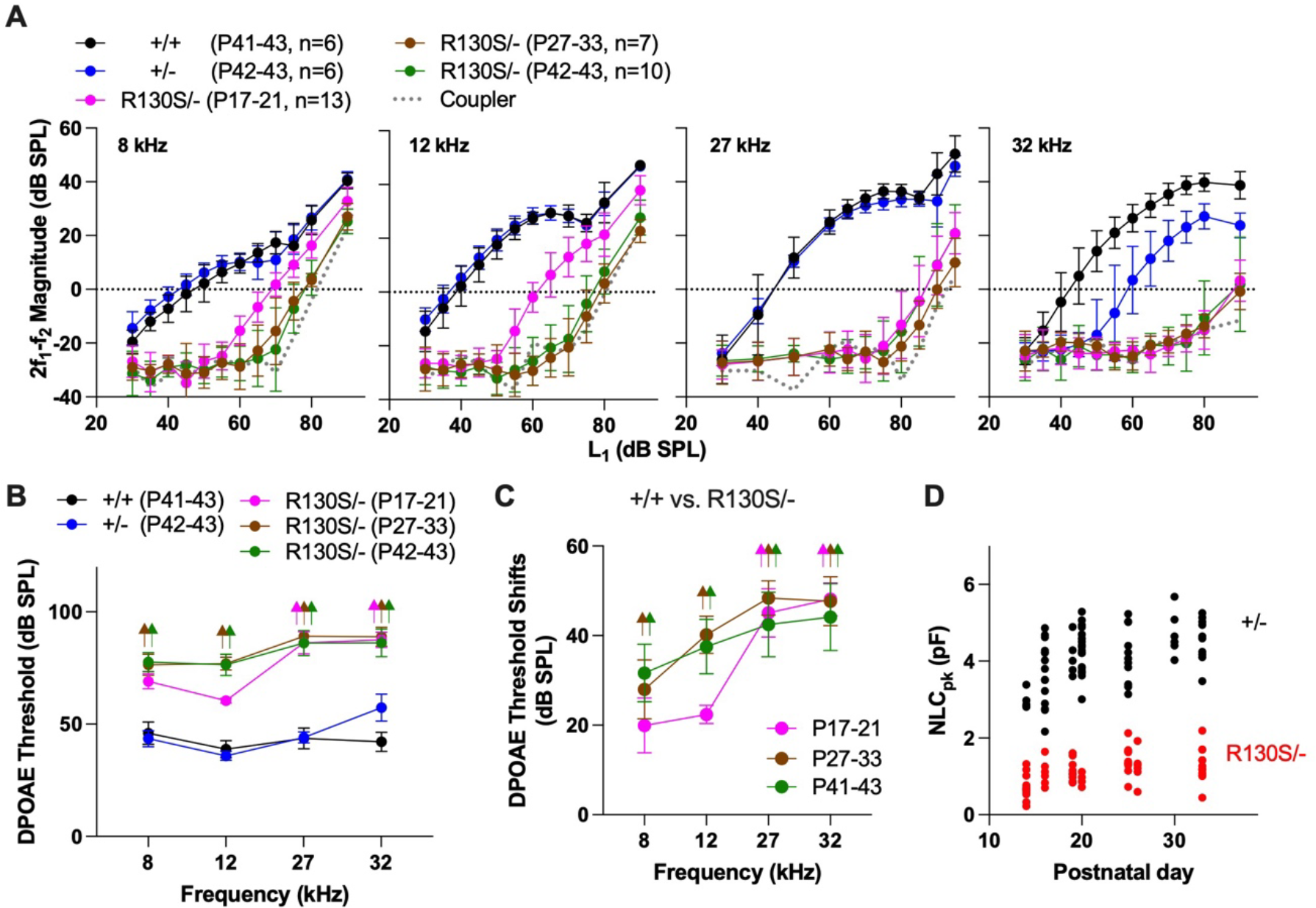
DPOAE input-output functions. (**A**) Growth functions were measured at f_2_ = 8, 12, 27, and 32 kHz (f_2_/f_1_=1.2, L_1_=L_2_+10 dB). Gray dotted lines in each panel indicate the coupler values, i.e., distortion in the sound. (**B**) DPOAE thresholds determined from panel A using age-matched controls. WT data for the two groups of younger controls were not included in Figs. 3 and 4A for clarity. Upward arrows for *Slc26a5*^*R130S/-*^ indicate thresholds that are similar to coupler values. Only in the very youngest mice at f_2_=8 and 12 kHz are thresholds independent of distortion in the stimulus. (**C**) DPOAE threshold shifts between *Slc26a5*^*R130S/-*^ vs. WT mice. Thresholds for WT mice at P20-21 and P27-33 are not provided in panel B. (**D**) The magnitudes of NLC at various ages between P14 and P33. NLC was recorded in isolated OHCs, and NLC_pk_ determined as in Fig. 2.

### Premature OHC loss in *Slc26a5*^*R130S/-*^ compound heterozygotes

Contribution of prestin to OHC maintenance is evidenced by the fact that prestin-KO mice suffer from premature and progressive OHC loss (Liberman et al., 2002; Wu et al., 2004). R130S-prestin, though not truncated, may not fully contribute to OHC maintenance, because its membrane targeting is partially impaired (Takahashi et al., 2016a; Takahashi et al., 2023). In fact, cytocochleograms of *Slc26a5*^*R130S/R130S*^ homozygous mice showed signs of OHC loss at high frequencies at six weeks of age (Takahashi et al., 2023). To examine the contribution of R130S-prestin to OHC maintenance in *Slc26a5*^*R130S/-*^ compound heterozygous mice, we constructed cytocochleograms at P21, P31, and at P42-43. We found minimal OHC loss at weaning. However, by six weeks of age the OHC loss becomes evident in cochleae from *Slc26a5*^*R130S/-*^ (**Figs. 5**) and exceeds that reported in *Slc26a5*^*R130S/R130S*^ (Takahashi et al., 2023). For the *Slc26a5*^*R130S/-*^ compound heterozygous mice, there are missing OHCs especially in regions coding for frequencies greater than 20 kHz. In contrast, *Slc26a5*^*R130S/R130S*^ homozygous mice retained more OHCs at these same locations (Takahashi et al., 2023).

**Fig. 5.**
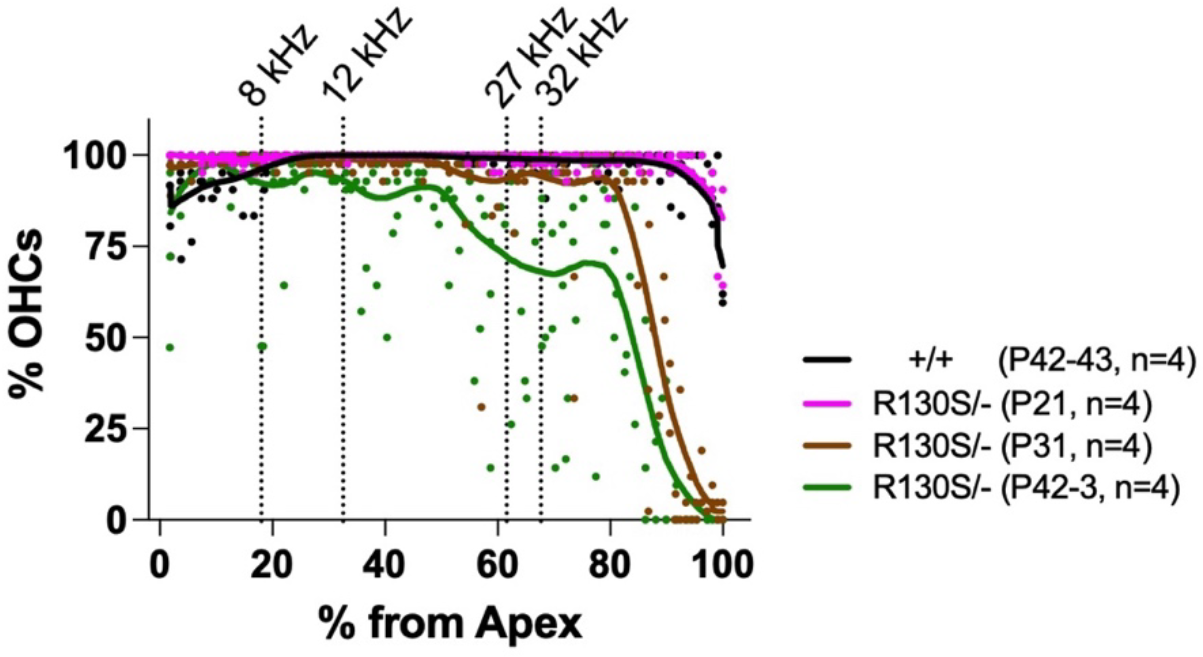
Progressive OHC loss in *Slc26a5*^*R130S/-*^ compound heterozygotes. Cytocochleograms of *Slc26a5*^*R130S/-*^ mice at P21, P31, and P42-43. Solid lines indicate LOWESS curve fits to better visualize the degree of OHC loss. Vertical dotted lines correspond to the f_2_ frequencies used for the DPOAE measurements in Fig. 4.

## DISCUSSION

The objective of this study was to fully characterize the pathophysiological consequences of the p.R130S prestin missense variant using *Scl26a5*^*R130S/-*^ compound heterozygous mice as a DFNB61 model for human *SLC26A5*^*W70X/R130S*^ compound heterozygous patients. As anticipated, we confirmed OHC dysfunction in this mouse model, which is most credibly ascribed to slowed motor kinetics of R130S-prestin as claimed in our recent study for *Slc26a5*^*R130S/R130S*^ mice (Takahashi et al., 2023). We also found that *Scl26a5*^*R130S/-*^ compound heterozygotes subsequently suffered progressive OHC degeneration, defining an additional pathogenic role of the p.R130S prestin variant in DFNB61 hearing loss.

Given the recessive nature of DFNB61-associated prestin variants and premature OHC loss found in prestin-KO mice, it is likely that loss-of-function underlies degeneration of OHCs expressing R130S-prestin. The partially impaired membrane targeting of R130S-prestin reported previously (Takahashi et al., 2016a; Takahashi et al., 2023), and further confirmed in this study, is in line with such a loss-of-function-based pathology. Since prestin belongs to the SLC26 anion transport family, it was speculated that OHC loss found in prestin-KO mice might be ascribed to the loss of prestin-mediated anion transport activity (Liberman et al., 2002). However, selective abrogation of prestin’s anion transport function did not affect OHC maintenance for at least seven months in S396D-prestin homozygous mice (Takahashi et al., 2023). It should also be noted that prestin hypomorphic mice do not suffer from OHC loss despite drastic reduction in WT prestin expression (Yamashita et al., 2012), suggesting that reducing prestin protein in the OHC lateral membrane *per se* would not necessarily lead to OHC degeneration.

One possible explanation for why OHC maintenance is differentially affected in various mutants may relate to a mismatch in mechanical impedance and how this relates to cell death-inducing mechanical stress. In the organ of Corti, OHCs are physically in contact with the tectorial membrane (TM) and with Deiters’ cells, thereby forming a tightly coupled feedback loop. The absence of prestin makes OHCs short and mechanically compliant (Cheatham et al., 2007; Dallos et al., 2008; Liberman et al., 2002; Wu et al., 2004), resulting in a mechanical impedance mismatch within the feedback loop that could contribute to OHC dysfunction in prestin-KO mice. Therefore, it is conceivable that reduced axial stiffness renders prestin-lacking OHCs more susceptible to sound-induced mechanical displacements. Progressive degeneration of R130S-prestin-expressing OHCs might also relate to reduced OHC stiffness that is ascribed to a decrease in the chloride binding affinity of R130S-prestin, since prestin-associated OHC axial stiffness depends on intracellular chloride (He et al., 2003).

Alternatively, it is possible that OHC degeneration may be ascribed to prestin’s slowed or absent motor activity, rather than pertaining to its presence as a static building block in the OHC’s lateral membrane. Sound-induced displacement of the basilar membrane (BM) towards the scala vestibuli deflects the stereocilia of OHCs in the excitatory direction and depolarizes the cells, resulting in somatic contraction of OHCs. This electromotility slightly enhances the BM motion towards scala vestibuli, while the TM moves in the opposite direction, i.e., to scala tympani. In other words, the top and bottom of the OHC move in opposite directions on a cycle-by-cycle basis in response to excitatory sinusoidal stimulation (Dewey et al., 2021). In addition, it is important to note that cochlear amplification augments fluid motion between the TM and the reticular lamina, which deflects the stereocilia of IHCs that are free standing (Dallos, 1985). In contrast, the tallest OHC stereocilia are firmly attached to the TM. Residence within this tightly coupled feedback loop may explain, at least in part, the OHCs vulnerability to sound-induced mechanical insults compared to IHCs. In this scenario, the reduction or elimination of OHC somatic contractions could enhance mechanical stress, especially in basal sensory receptors that are active even at rest (Beurg et al., 2010). This hypothetical protection requires prestin’s motor activity and must occur on a cycle-by-cycle basis. In the absence of prestin or when prestin’s motor kinetics are slowed, the maintenance of OHCs especially at the base of the cochlea is compromised.

## METHODS

### Stable HEK293T cell lines and immunofluorescence

cDNAs encoding WT and W70X-prestin were cloned into a pSBtet-pur vector (Addgene) using SfiI sites with or without a C-terminal mTurquoise (mTq2) tag. Stable cell lines expressing these prestin constructs in a doxycycline-dependent manner were established in HEK293T cells as previously described (Wasano et al., 2020). Briefly, pSB plasmids were introduced together with pCMV(CAT)T7-SB100 (Addgene) using Effectene (Qiagen), and the transfected cells were selected in DMEM supplemented with 10% FBS and 1μg/ml puromycin (Fisher Scientific). To examine WT and W70X-prestin expression, stable cells were seeded on a cover glass. After two days of 1 μg/ml doxycycline addition to the media, cells were fixed with 2% PFA in PBS for 10 minutes at room temperature, rinsed thrice with PBS, and mounted using DAKO fluorescent mounting medium (Agilent). For untagged prestin, anti-N-terminal prestin antisera (Zheng et al., 2005) was used followed by Alexa Fluor 546-conjugated anti-rabbit secondary antibody (Thermo A-11035). Images were captured using a Nikon A1R confocal microscope as previously described (Takahashi et al., 2019).

### Animals

A mouse model expressing p.R130S prestin on the FVB/NJ background was generated previously (Takahashi et al., 2023). Prestin-KO heterozygotes on FVB/NJ background (Cheatham et al., 2015) were crossed with *Slc26a5*^*R130S/+*^ heterozygotes to obtain *Slc26a5*^*R130S/-*^ compound heterozygotes along with WT and prestin-KO littermates. Genotyping was outsourced to Transnetyx (Cordova, TN). All procedures and protocols were approved by Northwestern University’s Institutional Animal Care and Use Committee (IACUC) and by NIDCD.

### DPOAE measurement

Distortion product otoacoustic emissions (DPOAEs) were recorded using a custom probe placed close to the eardrum as described previously (Cheatham et al., 2014). DPOAE iso-input functions were recorded using two pure tones with fixed frequency ratio, f_2_/f_1_=1.2, and at f_2_ frequencies ranging from 1.9 to 47.0 kHz. The sound levels of both tones were set to 70 dB SPL. DPOAE input-output functions were also collected at f_2_ frequencies equal to 8, 12, 27 and 32 kHz. Although f_2_/f_1_ remained fixed at 1.2, the levels of f_1_ were 10 dB greater than those at f_2_ (L_1_ = L_2_ + 10 dB SPL). DPOAE thresholds were defined as the level of f_1_ that produced a 2f_1_-f_2_ component of 0 dB SPL. Coupler measurements were also made to determine distortion in the sound. These measures indicated that 2f_1_-f_2_ reached 0 dB SPL at 82, 79, 91, and 90 dB for f_2_=8, 12, 27, and 32 kHz, respectively. In order to indicate that the DPOAE “threshold” was due to distortion in the sound and could not be used as an assay for OHC function, upward pointing arrows were used in **Fig. 4**. For these recordings, mice were anesthetized with intraperitoneal injection of ketamine (100 mg/kg) and xylazine (10 mg/kg) with supplemental injections given as needed. All emission measurements were from the animal’s left ear in a sound-isolated and electrically-shielded chamber. A heating pad was used to maintain body temperature at approximately 37°C.

### NLC recordings

Apical OHCs from a region of the cochlea corresponding to 4-10 kHz were isolated from mice as described previously (Homma et al., 2013). Whole-cell recordings (for OHCs and HEK293T cells) were performed at room temperature using the Axopatch 200B amplifier (Molecular Devices) with a 10 kHz low-pass filter. Recording pipettes pulled from borosilicate glass were filled with an intracellular solution containing (mM): 140 CsCl, 2 MgCl_2_, 10 EGTA, and 10 HEPES (pH 7.4). Cells were bathed in an extracellular solution containing (mM): 120 NaCl, 20 TEA-Cl, 2 CoCl_2_, 2 MgCl_2_, 10 HEPES (pH 7.4). Osmolality was adjusted to 309 mOsmol/kg with glucose. The initial bath resistances were 3-4 MΩ. NLC recordings were performed using a 0 mV holding potential and a sinusoidal voltage stimulus (2.5-Hz, 120-150 mV amplitude) superimposed with two higher frequency stimuli (390.6 and 781.2 Hz, 10 mV amplitude). While recording, the intracellular pressure was not adjusted. Data were collected and analyzed by jClamp (SciSoft Company, New Haven, CT) (Santos-Sacchi et al., 1998).

### NLC data analysis

Voltage-dependent C_m_ data were analyzed using the following two-state Boltzmann equation:

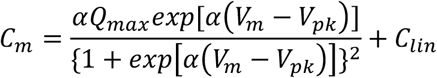

where α is the slope factor of the voltage-dependence of charge transfer, Q_max_ is the maximum charge transfer, V_m_ is the membrane potential, V_pk_ is the voltage at which the maximum charge movement is attained, and C_lin_ is the linear capacitance.

### Cytocochleograms

Mice were cardiac perfused with 4% paraformaldehyde and inner ears extracted. After fixation and subsequent decalcification, cochleae were dissected following the Eaton-Peabody Laboratory dissection protocol for whole mounts (Liberman et al., 2015). The organ of Corti was stained with anti-prestin (Zheng et al., 2005) followed by goat anti-rabbit Alexa Fluor 488 secondary antibody (ThermoFisher). Images were taken using a Leica DM IRB fluorescence microscope or Keyence BZ-X800 for OHC counts as previously described (Takahashi et al., 2016b). Cochlear locations corresponding to the frequencies tested for the DPOAE measurements were determined based on the mouse frequency-place map (Müller et al., 2005). LOWESS smooth fits (Cleveland, 1979) were included to facilitate visual inspection of the results (**Fig. 5**).

### Statistical analyses

Statistical analyses were performed using Prism (GraphPad Software). One-way ANOVA combined with the Tukey’s post hoc test was used for multiple comparisons. The uncertainties (σ) associated with subtraction computations to determine the differences in DPOAE thresholds with respect to WT controls (**Fig. 4C**) were calculated by the following equation:

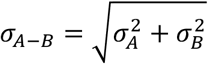

where A and B are the mean values with associated errors, σ_A_ and σ_B_, respectively.

## ACKNOWLEDGEMENTS

This work was supported by an NIH grant DC017482 (to KH) and by the Hugh Knowles Center. Some of the imaging work was performed at the Northwestern University Center for Advanced Microscopy generously supported by NCI CCSG P30 CA060553 awarded to the Robert H Lurie Comprehensive Cancer Center.

## Notes

**Competing Interest Statement:** The authors declare no competing financial interests or conflicts of interest.

### Competing Interest Statement

The authors have declared no competing interest.

